# Complex roles for reactive astrocytes in the triple transgenic mouse model of Alzheimer disease

**DOI:** 10.1101/797662

**Authors:** Océane Guillemaud, Kelly Ceyzériat, Thomas Saint-Georges, Karine Cambon, Fanny Petit, Lucile Ben Haim, Maria-Angeles Carrillo-de Sauvage, Martine Guillermier, Sueva Bernier, Anne-Sophie Hérard, Charlène Joséphine, Alexis Pierre Bémelmans, Emmanuel Brouillet, Philippe Hantraye, Gilles Bonvento, Carole Escartin

## Abstract

In Alzheimer disease (AD), astrocytes undergo complex changes and become reactive. The consequences of this reaction are still unclear. To evaluate the net impact of reactive astrocytes in AD, we recently developed viral vectors targeting astrocytes that either activate or inhibit the JAK2-STAT3 pathway, a central cascade controlling astrocyte reaction.

We aimed to evaluate whether reactive astrocytes contribute to Tau as well as amyloid pathologies in the hippocampus of 3xTg-AD mice, an AD model that develops Tau hyperphosphorylation and aggregation in addition to amyloid deposition. JAK2-STAT3 pathway-mediated modulation of reactive astrocytes in the hippocampus of 3xTg-AD mice, did not significantly influence Tau phosphorylation or amyloid processing and deposition, at early, advanced and terminal stage of the disease. Interestingly, inhibition of the JAK2-STAT3 pathway in hippocampal astrocytes did not improve short-term spatial memory in the Y maze but it reduced anxiety in the elevated plus maze. Our unique approach to specifically manipulate reactive astrocytes in situ show these cells may impact behavioral outcomes without influencing Tau or amyloid pathology.

## 1. INTRODUCTION

Alzheimer disease (AD) is a devastating neurodegenerative disease and the most common form of dementia (Querfurth and LaFerla, 2010). It is characterized by extracellular accumulation of amyloid plaques, Tau hyper-phosphorylation, synaptic alterations and neuronal degeneration. The exact mechanisms responsible for AD are still disputed (Sala Frigerio and De Strooper, 2016). The amyloid cascade hypothesis was first put forward. According to this hypothesis, successive cleavages of the amyloid precursor protein (APP) by β- and γ-secretases, generate high levels of soluble Aβ peptides that oligomerize and form amyloid plaques in the brain of AD patients. Aβ peptides cause synaptic dysfunction, neurodegeneration and cognitive impairment (Hardy and Higgins, 1992). This hypothesis was later refined to include Tau hyper-phosphorylation as a key pathogenic element (Kametani and Hasegawa, 2018; Medina and Avila, 2014). Hyperphosphorylated Tau proteins also oligomerize and form neurofibrillary tangles that inhibit microtubule assembly and alter neuronal functions. Progression of Tau pathology from entorhinal cortex to other isocortex regions (Braak and Braak, 1991) correlates better than amyloid pathology with cognitive impairment observed in patients (Arriagada et al., 1992; Masters et al., 2015). More recently, evidence emerged showing that AD is not only caused by cell-autonomous processes within neurons, and that glial cells may also play an instrumental role in disease progression (Arranz and De Strooper, 2019; De Strooper and Karran, 2016; Heneka et al., 2015).

In particular, astrocytes are in charge of key brain functions, including neurotransmitter recycling, ion homeostasis, metabolic and trophic support (Verkhratsky and Nedergaard, 2018). In the brain of AD patients or animal models, astrocytes are reactive and participate in neuroinflammation. Reactive astrocytes are classically characterized by morphological changes (hypertrophy and process reorganization), as well as upregulation of intermediate filament proteins [Glial fibrillary acidic protein (GFAP), vimentin, (Hol and Pekny, 2015)]. Much less is known about their functional features and how exactly they impact AD outcomes (Ben Haim et al., 2015a). Several studies have reported alterations in some of the multiple functions operated by astrocytes like glutamate uptake (Masliah et al., 1996), metabolic supply (Allaman et al., 2010; Sancheti et al., 2014), production of antioxidant molecules (Abeti et al., 2011; Allaman et al., 2010) or regulation of synaptic transmission (Delekate et al., 2014; Jo et al., 2014; Lian et al., 2015; Wu et al., 2014); for review see (Ben Haim et al., 2015a; Chun and Lee, 2018). Interestingly, astrocytes may directly participate in APP metabolism and Aβ clearance through various mechanisms, ranging from extracellular cleavage to phagocytosis or drainage through the glymphatic system (Ries and Sastre, 2016). But how these clearance mechanisms are altered when astrocytes acquire reactive features during AD progression is unknown. Similarly, the exact impact of reactive astrocytes on Tau hyperphosphorylation and aggregation remains to be established (Kahlson and Colodner, 2015; Leyns and Holtzman, 2017).

We recently developed a new method that efficiently blocks morphological and molecular changes in reactive astrocytes, by inhibiting the Janus kinase 2-Signal transducer and activator of transcription 3 (JAK2-STAT3) pathway, through astrocyte-specific expression of its inhibitor suppressor of cytokine signaling 3 [SOCS3, (Ceyzériat et al., 2018)]. We showed that reactive astrocytes promote amyloid deposition and learning deficits in the APP/PS1dE9 transgenic mouse model of AD, and early synaptic deficits in 3xTg-AD mice, another transgenic model that also develops Tau pathology. Here, we aimed to evaluate in 3xTg-AD mice, whether reactive astrocytes 1) influence Tau and amyloid pathologies at different disease stages and 2) have long-term effects on cognitive functions. Une
xpectedly, we found that JAK2-STAT3 activation in reactive astrocytes does not affect Tau or amyloid pathology in 3xTg-AD mice at early, advanced and terminal disease stages. In addition, inhibition of reactive astrocytes does not improve working memory but it reduces anxiety in 3xTg-AD mice.

## 2. MATERIEL AND METHODS

### Animals

Triple transgenic AD (here referred to as 3xTg mice) mice express human APPswe and human Tau^P301L^ under a Thy-1 promoter as well as a point mutation on the mouse *Psen1* gene (PS1^M146V^), on a mixed C57BL/6J × 129Sv background (Oddo et al., 2003). C57BL/6J × 129Sv mice were used as controls. Only females were used in this study, as they display earlier neuropathology (Carroll et al., 2010; Hirata-Fukae et al., 2008). Breeding pairs were obtained from the Mutant Mouse Regional Resource Centers. All experimental protocols were approved by a local ethics committee (CETEA N°44) and submitted to the French Ministry of Education and Research (Approvals # APAFIS#4565-20 16031711426915 v3 and APAFIS#4503-2016031409023019). They were performed in an authorized facility (#D92-032-02), under the supervision of a veterinarian and in strict accordance with the recommendations of the European Union (2010-63/EEC) for the care and use of laboratory animals. All efforts were made to promote animal welfare. Mice were housed under standard environmental conditions (12-hour light-dark cycle, temperature: 22 ± 1°C and humidity: 50%) with *ad libitum* access to food and water. Mice of the appropriate genotype were randomly allocated to experimental groups.

### Stereotactic injections of viral vectors

Adeno-associated viruses of serotype 9 (AAV2/9) encoding *Gfp*, murine *Socs3*, or a constitutive active form of *Jak2* (JAK2^T875N^ or JAK2ca) were produced and tittered at the MIRCen viral vector facility, as described previously (Ceyzériat et al., 2018). All transgenes were under the control of the gfaABC1D promoter, which drives specific expression in hippocampal astrocytes (Ceyzériat et al., 2018). Before surgery, mice were anesthetized with a mixture of ketamine (100 mg/kg) and xylazine (10 mg/kg) and received *s.c.* injection of lidocaine (7 mg/kg) at the incision site, 10 min before injection. Mice received paracetamol in drinking water (1.6 mg/ml) for 48 h after surgery. AAV were diluted in 0.1 M phosphate buffer saline (PBS) with 0.001% pluronic acid, at a final total concentration of 2.5 10^9^ viral genome/μl. Mice were positioned on a stereotactic apparatus with tooth bar set at 0 mm. They were injected in the CA1 region of the hippocampus (coordinates: - 3 mm antero-posterior, +/− 3 mm lateral and – 1.5 mm ventral, from dura) with 2 μl of AAV suspensions at a rate of 0.2 μl/min. After 5 min, the cannula was slowly removed and the skin sutured. Mice recovered for at least 2 months before analysis.

### Experimental groups

Four mouse cohorts were analyzed (**Supplemental Fig. 1**). The number of mice analyzed is mentioned in each figure.

1. Homozygous 3xTg mice were injected at 6 months with AAV-GFP (3xTg-GFP mice) or AAV-SOCS3 + AAV-GFP at the same total viral titer (3xTg-SOCS3 mice), to block reactive astrocytes. AAV-GFP was co-injected with AAV-SOCS3 to visualize infected cells. WT mice of the same age, gender and genetic background were injected with AAV-GFP (WT-GFP) as control. Mice were tested on the elevated plus maze and the Y maze at 8 and 14-15 months. Mice were euthanized by an overdose of pentobarbital at the age of 16 months. The two brain hemispheres were rapidly dissected on ice. One was drop-fixed in 4% paraformaldehyde (PFA) and used for immunohistochemistry. The other was cut into 1-mm-thick slices on ice and the hippocampal GFP^+^ area was dissected out under a fluorescent macroscope, snap frozen in liquid nitrogen and stored at −80°C until protein extraction.
2. At 3 months, WT mice were injected with AAV-GFP, and 3xTg mice were injected with AAV-GFP or with AAV-JAK2ca + AAV-GFP (same total viral titer, 3xTg-JAK2ca mice), to exacerbate astrocyte reaction. Mice were euthanized as 16 months and analyzed by histological and biochemical approaches as described for the first cohort.
3. At 4 months, WT mice were injected with AAV-GFP, and 3xTg mice were injected with AAV-GFP or with AAV-SOCS3 + AAV-GFP. Mice were euthanized by an overdose of pentobarbital at the age of 9 months. The hippocampus was dissected out, snap frozen in liquid nitrogen and stored at −80°C until protein extraction.
4. At 24 months, WT mice were injected with AAV-GFP, and 3xTg mice were injected with AAV-GFP or with AAV-SOCS3 + AAV-GFP. Mice were analyzed by biochemical approaches at 26 months as described for cohort 3.

### Immunohistology

Dedicated hemispheres were post-fixed for 24 h in 4% PFA, cryoprotected in a 30% sucrose solution and cut on a freezing microtome into serial 30-μm-thick coronal sections. Series were stored at −20°C in an anti-freeze solution until used for immunostaining.

#### Immunofluorescence

Sections were rinsed in PBS for 3 × 10 min and blocked in 4.5% normal goat serum (NGS) diluted in PBS with 0.2% Triton X-100 (PBST), for 1h at room temperature (RT). Sections were incubated overnight at 4°C with anti-GFAP-Cy3 antibody (1:1,000, Sigma, #C9205, Saint Louis, MO) diluted in 3% NGS/PBST. Sections were rinsed 3 × 10 min in PBS and incubated with secondary Alexa Fluor-conjugated antibodies (1:1,000, Invitrogen, Carlsbad, CA) in 3% NGS/PBST for 1h at RT. After 3 washes in PBS, sections were incubated overnight at 4°C with an anti-GFP biotinylated antibody (1:500, Vector Laboratories, #BA-0702, Burlingame, CA) in 3% NGS/PBST. After 3 rinses in PBS, sections were incubated for 1h at RT with Streptavidine-FITC (1:1,000, ThermoFisher Scientific, Waltham, MA) in 3% NGS/PBST and rinsed 3 times with PBS before being mounted on SuperFrost® Plus slides (ThermoFisher Scientific) and coverslipped with Fluorsave™ medium (Calbiochem, Darmstadt, Germany). For Iba1 (1:500, Wako, #191974, Neuss, Germany) and Vimentin (1:750, Abcam, #Ab24525, Cambridge) staining, a standard protocol was performed as described above but endogenous GFP was not amplified and sections were incubated with DAPI (1,2000, Invitrogen, #D21490) before being coverslipped with Vectashield™ medium (Vector Laboratories, #H1400). For methoxy-XO4 (MXO4, Tocris, Minneapolis, MN) staining, sections were incubated with 33 μg/ml MXO4 in 0.1 M PBS, for 30 min at RT under mild agitation. After 3 rinses with PBS, a standard protocol for immunofluorescence was performed as described above. For Aβ staining with the 4G8 antibody (1:500, Covance, #SIG39240, Princeton, NJ), sections were treated with 70% formic acid during 2 min before incubation with primary biotinylated 4G8 antibody for 48h at 4°C. After 3 rinses with PBS, sections were incubated for 1h at RT with Streptavidine-Cy3 (1:1,000, Sigma) in 3.5% NGS/PBST. Sections were then mounted as previously described.

#### Immunohistochemistry

Sections were rinsed in PBS for 3 × 10 min and incubated with H_2_O_2_ (1:100, Sigma) for 20 min at RT to inhibit peroxidases. For HT7 antibody, sections were pre-treated with 10 mM citrate buffer for 20 min at 90°C before H_2_0_2_ treatment. Sections were then blocked in 4.5% NGS/PBST for 1h at RT and incubated overnight at RT or 48h at 4°C in 3% NGS/PBST with the following primary antibodies: anti-Aβ42 (1:500, Rabbit, Life Technologies, #44-344, Carlsbad, CA), AT8 (1:400, Mouse, ThermoFisher Scientific, #MN1020B), HT7 (1:1,000, Mouse, Innogenetics, #90204, Zwijnaarde, Belgium), anti-phospho-Ser422-Tau (1:400, Rabbit, Abcam, #Ab79415, Cambridge, UK). After 3 × 10 min rinses in PBS, sections were incubated for 1h at RT with biotinylated secondary antibodies (1:4,000, except at 1:1,000 for anti-phospho-Ser422-Tau, Vector Laboratories) in 3% NGS/PBST. They were washed 3 × 10 min and incubated for 1h at RT with avidin/biotin complexes (1:250 in PBST, Vector Laboratories) and after 3 washes with PBS, incubated with VIP (Vector Laboratories). Sections were mounted on SuperFrost Plus slides and dried overnight at RT. Sections were dehydrated in acetone and xylene and coverslipped with Eukitt mounting medium (Chem-Lab, Zedelmgem, Belgium).

### Immunostaining quantification

GFAP and Vimentin stainings were quantified on 10×-tiled images acquired with an epifluorescence microscope (BM6000B, Leica, Nussloch, Germany). Virally transduced GFP^+^ area was manually segmented and the GFAP and Vimentin mean grey values were measured in this GFP^+^ region with Image J. Representative fluorescent images in Fig. 1 were acquired with a 40× objective on a confocal microscope (TCS SP8, Leica).

**Figure 1.**
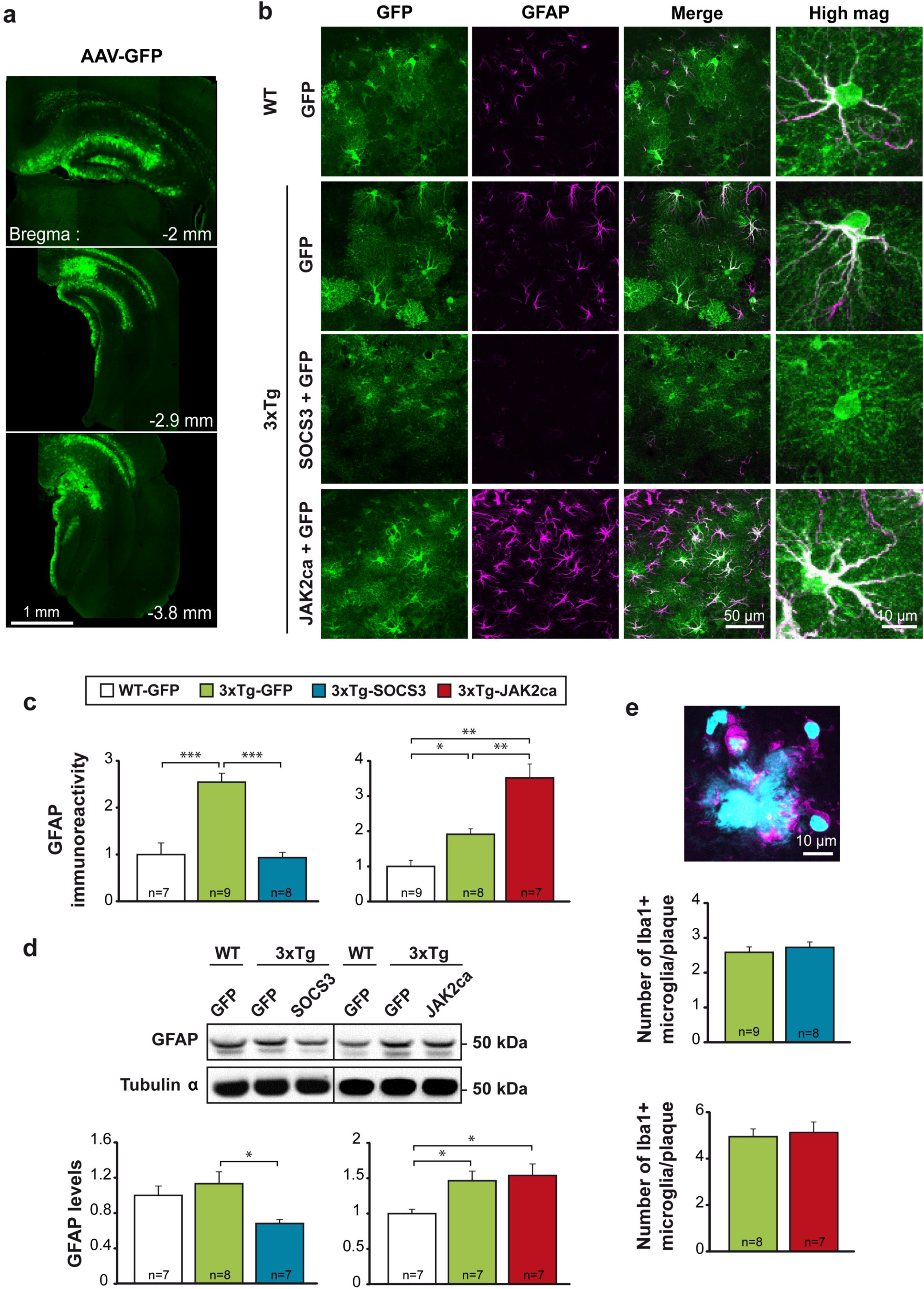
Long-term modulation of reactive astrocytes by targeting the JAK2-STAT3 pathway in 3xTg mice. **a**, Low magnification image of serial brain sections of a mouse injected with an AAV-GFP in CA1. GFP (green) is expressed throughout the dorsal hippocampus. **b**, Confocal images of GFP^+^astrocytes (green) stained for GFAP (magenta) in 16-month-old WT and 3xTg mice injected in the CA1 region with AAV-GFP, AAV-SOCS3 + AAV-GFP or AAV-JAK2ca + AAV-GFP (same viral titer). In 3xTg mice, astrocytes display the typical reactive features (GFAP overexpression and morphological changes). SOCS3 significantly reduces GFAP levels, while JAK2ca increases them. **c**, Quantification of GFAP immunoreactivity in the hippocampus. * *p* < 0.05, ** *p* < 0.01, *** *p* < 0.001. Kruskall-Wallis and Mann-Whitney tests. **d**, Western blotting and quantification of GFAP levels (normalized to tubulin α) in WT-GFP, 3xTg-GFP, 3xTg-SOCS3 and 3xTg-JAK2ca mice show the same pattern of GFAP modulation by SOCS3 and JAK2ca. * *p* < 0.05. One-way ANOVA and Tukey’s post hoc test. Quantifications are expressed relatively to the WT group in each cohort, whose value was set to 1. **e**, Representative confocal image of Iba1^+^ microglia cells (magenta) around an amyloid plaque (cyan) in 16-month-old 3xTg mice. Nuclei and amyloid plaques are labelled with DAPI (cyan). Quantification of the number of Iba1^+^ cells around a plaque shows no difference between groups. Student *t* test.

The number of Iba1 positive cells around plaque was quantified directly by scanning the whole slice on the z axis, under the 40× objective of the confocal microscope. An image was acquired at the center of each plaque to measure its cross area.

Serial sections stained for HT7, Phospho-Ser422-Tau and AT8 were scanned under bright light, with an Axio scanZ.1 (Zeiss, Oberkochen, Germany). The immune-positive area within the manually-segmented hippocampus was automatically detected on each section, with an intensity threshold applied with ImageJ. The immunopositive volume was calculated by the Cavalieri method, normalized to the total volume of the hippocampus (which was not different between groups), and expressed as a percentage. The number and surface of MXO4-labelled plaques was quantified on 10×-tiled fluorescent images. Aβ42 plaques were analyzed on immunostained sections scanned under bright field on the AxioSan. Automatic detection of objects with intensity and size thresholds was performed on these images, after manual segmentation of the hippocampus on each section, with Image J. To quantify 4G8 immunoreactivity, the labeled area in the *subiculum* and *stratum pyramidale* of the CA1 region, was manually segmented on 10×-tiled fluorescent images and the mean grey value was measured with Image J.

### Protein extraction

Hippocampal samples were homogenized by sonication (Labortechnik, Wasserburg, Germany, 6 strokes with cycle = 0.5, 30% amplitude) in lysis buffer [50 mM Tris-HCl pH = 7.4, 150 mM NaCl, 1% Triton X-100 with 1:100 phosphatase inhibitors (Sigma, cocktail 2) and 1× protease inhibitors (Roche, Basel, Switzerland)] centrifuged at 20,000 g for 20 min at 4°C. The supernatant contains Triton X-100-soluble proteins and was used for western blotting and MSD® ELISA tests. Protein concentration was measured by the BCA test (Pierce, Waltham, MA).

### Western blot

Samples were diluted in loading buffer with DTT (NuPAGE® LDS sample buffer and sample reducing agent, Invitrogen). Ten μg of proteins were loaded on a NuPAGE™ 4-12% Bis-Tris Midi Gel (Life Technologies) and exposed to 200 V for 45 min in NuPAGE™ Running Buffer (Invitrogen). Proteins were transferred on a nitrocellulose membrane with an iBlot Gel transfer device (Invitrogen). After 3 × 10 min rinses in Tris buffer saline and 0.1% Tween 20 (TBST), membranes were blocked in 5% milk in TBST for 1h at RT and incubated for 3h at RT, or overnight at 4°C with the following primary antibodies: 6E10 (1:500, Mouse, Covance, #SIG-39320-20), anti-ApoE (1:1,000, Rabbit, Abcam, #Ab20874), anti-BACE1 (1:1,000, Rabbit, Cell signaling, #5606P, Danvers, MA), anti-GFAP (1:5,000, Rabbit, Dako, #Z0334, Troy, MI), anti-IDE (1:400, Rabbit, Abcam, #Ab32216), anti phospho-Ser404-Tau (1:1000, Rabbit, Sigma, #T7444) and anti-Tubulin α (1:1,000, Mouse, Sigma, #T5168). After 3 × 10 min washes in TBST, membranes were incubated for 1h at RT with HRP-conjugated secondary antibodies (1:5,000, Vector laboratories) diluted in TBST with 5% milk. Membranes were incubated with the Clarity Western ECL substrate (Bio-Rad, Hercules, CA) and the signal was detected with a Fusion FX7 camera (ThermoFisher Scientific). Band intensity was quantified with ImageJ and normalized to Tubulin α. Each antibody was used on at least 2 different membranes. Representative images and quantification are shown.

### MSD® ELISA tests

Triton x-100 soluble proteins were diluted in the diluent provided for the V-PLEX Aβ peptide panel kit (Aβ38, Aβ40, Aβ42, 6E10 antibody, MSD®, Rockville, MD). For the phospho-Thr231-Tau/Total Tau kit (MSD®), samples were diluted in 20 mM Tris HCl pH = 7.4, 150 mM NaCl, 1 mM EDTA, 1 mM EGTA, 1% Triton X-100, with phosphatase and protease inhibitors. For both assays, samples were loaded in triplicate and manufacturer’s protocol was followed. Peptide levels were quantified with a standard curve, using the Discovery Workbench4.0, MSD® software, and normalized to the protein content in each well.

### Behavioral analysis

WT-GFP, 3xTg-GFP and 3xTg-SOCS3 mice were tested at the age of 8 and 14-15 months on the elevated plus maze to evaluate anxiety and after one month, on the Y maze to assess working spatial memory. All tests were performed between 8 am and noon. Mice were handled daily for 2 min for 5 d before the first training session. Both mazes were cleaned with 10% ethanol and cautiously dried between each phase and each mouse to remove odors.

The elevated plus maze test was formed by four connected arms placed 50 cm above the ground, illuminated at 100 lux in the center (see **Fig. 5b**). Two arms made of black Plexiglas were closed (non-stressful arms) while the two others were open (stressful arms). Each mouse was placed in the center of the maze and video-tracked for 5 min with the EthoVision XT11.5 software (Noldus IT, The Netherlands). The following parameters were measured: distance traveled and time spent in each arm, frequency of entry into each arm, duration and number of head dipping (counted manually during the experiment). Mice performing fewer than 4 entries into different arms were excluded from the analysis.

The Y maze test was composed of three arms in black Plexiglas, illuminated at 110 lux in the center. Each mouse was placed at the extremity of the “start arm” and left for a training session of 15 min, with one of the two other arms closed with a black Plexiglas door (the closed arm was randomly chosen for each mouse). Then, the mouse was brought back to its cage for 5 min, the maze was cleaned, the closed arm was opened and the mouse placed at the extremity of the start arm for a 5 min test session, under videotracking (see **Fig. 5a**). The distance traveled and time spent in each arm (start, familiar, and novel arms) were measured and analyzed with EthoVision. Given their natural curiosity, mice are expected to spend more time exploring the new arm. Mice that stayed in the start arm during training or made less than 4 total entries in the 3 arms during testing were excluded.

### Statistical analysis

Results are expressed as mean ± SEM. Statistical analysis were performed with Statistica software (StatSoft, Tulsa, OK). Student *t* test and one-way ANOVA and Tukey’s post hoc test were used to compare two and three groups, respectively. Two-way (group, arm) ANOVA followed by Tuckey’s post-hoc test was used for Y maze analysis. For each analysis, normality of variables or residues and homoscedasticity were assessed. If any or both conditions of application were not fulfilled, non-parametric tests were used. Two groups were compared by the Mann-Whitney test and 3 independent groups were compared by a Kruskal-Wallis test followed by a Mann-Whitney test. Investigators were partially blinded to the group, when performing experiments and measuring outcomes (as the group can be guessed based on the presence of amyloid or Tau staining or GFP levels for example). The significance level was set at *p* < 0.05. The number of analyzed mice per group is indicated in each bar histogram.

## 3. RESULTS

### 3.1 The JAK2-STAT3 pathway efficiently controls reactive features of hippocampal astrocytes in 3xTg mice

We previously showed that adeno-associated viral (AAV) gene transfer of SOCS3, an inhibitor of the JAK-STAT3 pathway, in hippocampal astrocytes, prevents their acquisition of reactive molecular features, including in 9 and 12-month-old 3xTg mice (Ben Haim et al., 2015b; Ceyzériat et al., 2018). We studied whether SOCS3 was still able to block reactive astrocytes at a later stage, when both Tau and amyloid pathologies are present in 16-month-old 3xTg mice.

Six-month-old female 3xTg mice were injected with an AAV2/9 vector encoding *Socs3* under the gfaABC_1_D promoter (3xTg-SOCS3 mice). This viral vector allows the selective transduction of astrocytes, in the dorsal hippocampus (**Fig. 1a**), covering 25% of the whole hippocampus. WT and 3xTg mice were injected with an AAV encoding GFP as controls (WT-GFP and 3xTg-GFP mice respectively, see **Supplemental Fig. 1**). Brain sections from 16-month-old 3xTg-GFP mice displayed 3-fold higher GFAP levels than WT-GFP mice (**Fig. 1b, c**). In 3xTg-GFP mice, hippocampal astrocytes appear hypertrophic, a typical hallmark of reactive astrocytes. AAV-gene transfer of SOCS3 in astrocytes reduced GFAP expression and normalized astrocyte morphology (**Fig. 1b, c**). Mirror experiments were performed on another mouse cohort to further increase the activation of astrocytes by stimulation of the JAK2-STAT3 pathway. Three-month-old female 3xTg mice were injected in the hippocampus with an AAV encoding a constitutively active form of JAK2 (JAK2ca), the upstream kinase of the JAK2-STAT3 pathway, and were studied at 16 months (3xTg-JAK2ca mice). Control groups included WT-GFP and 3xTg-GFP mice injected and analyzed at the same age (**Supplemental Fig. 1**). 3xTg-JAK2ca mice displayed enhanced astrocyte reaction, visible as a two-fold increase in GFAP immunoreactivity and exacerbated enlargement and tortuosity of astrocyte processes, compared with 3xTg-GFP mice (**Fig. 1b, c**). Modulation of GFAP levels by SOCS3 and JAK2ca was also observed by western blotting on hippocampal samples taken from WT-GFP, 3xTg-GFP, 3xTg-SOCS3 and 3xTg-JAK2ca mice (**Fig. 1d**). The immunoreactivity for vimentin, another intermediate filament characteristic of reactive astrocytes was also reduced by more than 50% by SOCS3 in the hippocampus of 3xTg mice, although it did not reach significance (*p* = 0.065, Mann Whitney test). Conversely, vimentin immunoreactivity was increased more than 3-fold by JAK2ca (*p* = 0.018, Mann Whitney test). The number of microglia cells in direct contact with amyloid plaques was not impacted by either SOCS3 or JAK2ca in 3xTg-AD mice (**Fig. 1e**), suggesting that our strategy only affects astrocytes, as reported in another mouse model of AD (Ceyzériat et al., 2018).

These results show that manipulation of the JAK2-STAT3 pathway in hippocampal astrocytes is an efficient strategy to modulate the reactive state of astrocytes in 3xTg mice. We next evaluated how manipulation of reactive astrocytes by SOCS3 and JAK2ca impacted Tau hyperphosphorylation and amyloid deposition, two cardinal features of AD.

### 3.2 Modulation of reactive astrocytes does not influence Tau phosphorylation in 3xTg mice

At the age of 16 months, 3xTg mice display both Tau and amyloid pathologies. Tau can be phosphorylated on multiple epitopes. Three main approaches were used to characterize the effects of reactive astrocytes on Tau phosphorylation: i) immunostainings with antibodies against phospho-Ser422-Tau, phospho-Ser202/Thr205-Tau (AT8 antibody) and total human Tau (HT7 antibody) as a control, ii) ELISA MSD® assay with antibodies against phospho-Thr231-Tau (AT180 antibody) and total Tau and iii) western blotting against phospho-Ser404-Tau.

Positive staining for HT7, phospho-Ser422-Tau and AT8 was observed in 3xTg mice but not in WT mice. HT7 immunoreactivity was not changed by SOCS3 or JAK2ca in the hippocampus of 3xTg mice (**Fig. 2a, b**), suggesting that the expression of the human Tau transgene is not impacted by our experimental manipulations. Immunostaining with phospho-Ser422-Tau and AT8 antibodies revealed dystrophic neurons and numerous ghost tangles in the hippocampus (**Fig. 2a**). Immunoreactivity for these two antibodies was similar between the 3xTg-GFP, 3xTg-SOCS3 and 3xTg-JAK2ca groups (**Fig. 2b**), suggesting that modulation of reactive astrocytes through the JAK2-STAT3 pathway does not impact the level of phosphorylation of these epitopes.

**Figure 2.**
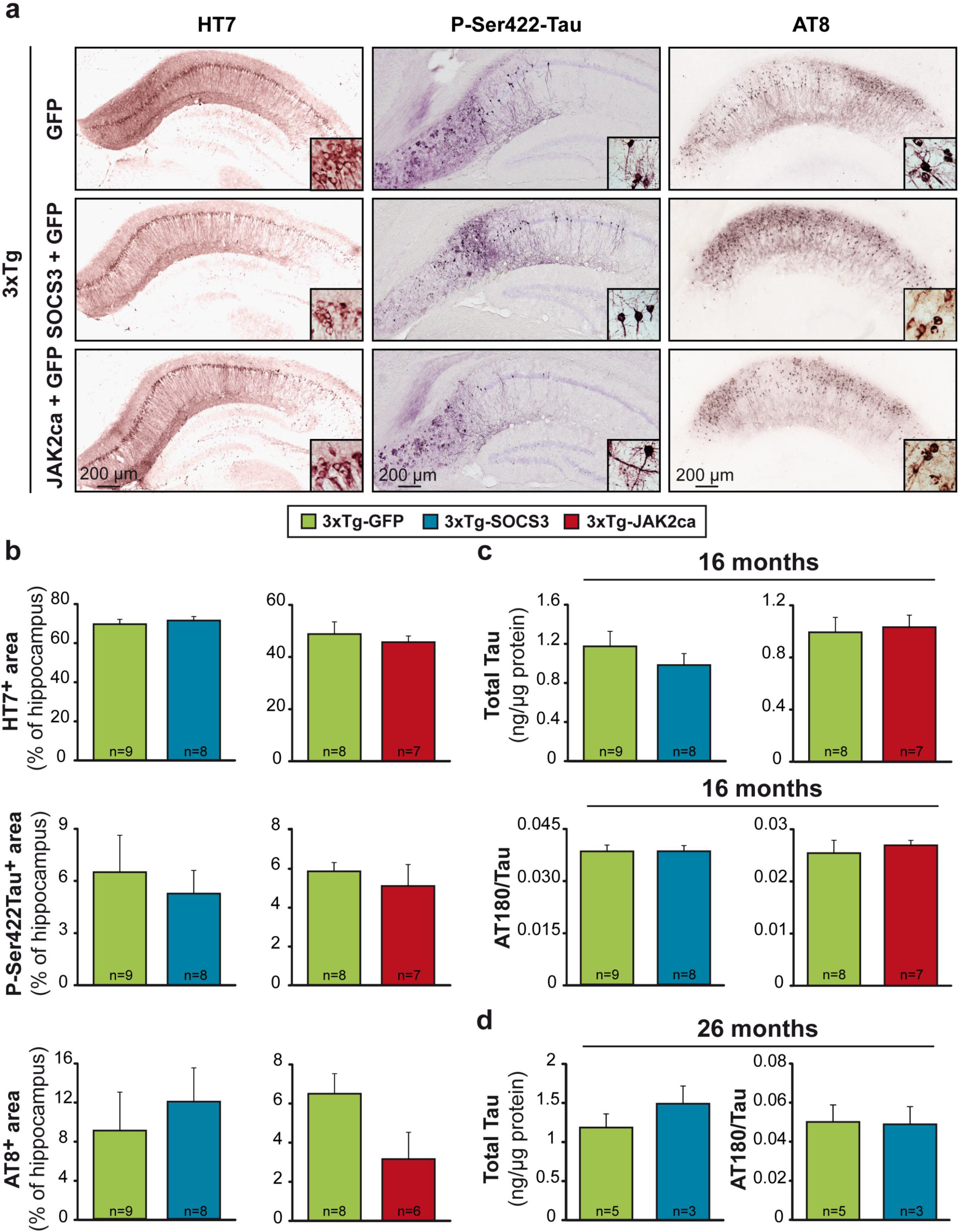
Modulation of reactive astrocytes does not impact Tau pathology in 3xTg mice. **a**, Representative brain sections of 16-month-old 3xTg-GFP, 3xTg-SOCS3 and 3xTg-JAK2ca mice stained with antibodies against total human Tau (HT7), phospho-Ser422-Tau (P-Ser422-Tau) or phospho-Ser202/Thr205-Tau (AT8). High magnification images for each staining shows dystrophic neurons and ghost tangles. **b**, Quantification of the immunopositive area in the whole hippocampus evidences no difference between groups for these 3 markers. Student *t* test (HT7 and AT8) or Mann-Whitney test (phospho-Ser422-Tau). **c**, MSD® measurement of phospho-Thr231-Tau (AT180 antibody) and total Tau in protein homogenates prepared from the hippocampus of 16-month-old 3xTg-GFP, 3xTg-SOCS3 and 3xTg-JAK2ca mice. Total Tau and AT180/Tau levels are not impacted by SOCS3 or JAK2ca in 3xTg mice. Mann-Whitney test. **d**, The same measurement in 26-month-old 3xTg-GFP and 3xTg-SOCS3 mice shows that total Tau and AT180/Tau levels are similar in both groups. Mann-Whitney test.

We next measured hippocampal concentrations of phospho-Thr231-Tau and total human Tau by MSD® assays on protein homogenates prepared from the hippocampus of 3xTg-GFP, 3xTg-SOCS3 or 3xTg-JAK2ca. Levels of phospho-Thr231-Tau normalized by total Tau and total Tau were not impacted by SOCS3 or JAK2ca in 16-month-old 3xTg mice (**Fig. 2c**). To evaluate whether reactive astrocytes impact Tau phosphorylation at a later stage of the disease, we performed the same measurement in 26-month-old female 3xTg mice that were injected 2 months before with AAV-GFP or AAV-SOCS3. Tau phosphorylation on Thr231 was also unaffected by SOCS3 in 26-month-old 3xTg mice (**Fig. 2d**).

Last, we analyzed phospho-Ser404-Tau levels by western blotting on hippocampal protein samples from 16-month-old 3xTg-GFP, 3xTg-SOCS3 and 3xTg-JAK2ca. We found no difference between groups (**Supplemental Fig. 2**).

Our results show that JAK2-STAT3 pathway modulation of reactive astrocytes in 3xTg mice does not influence Tau phosphorylation on several epitopes, at advanced and terminal stages of the disease.

### 3.3. Modulation of reactive astrocytes does not impact amyloid deposition in 3xTg mice

Amyloid pathology was assessed by four approaches: i) fluorescent staining with the Congo-red derivative Methoxy-XO4 (MXO4) that binds dense-core amyloid plaques of different size, ii) immunostaining with the Aβ42 antibody that labels small dense-core plaques (Serrano-Pozo et al., 2011) iii) immunofluorescent staining with the 4G8 antibody that labels diffuse plaques, as well as intracellular Aβ, which is prominent in 3xTg mice (Oddo et al., 2003), and finally iv) MSD® measurement of Triton X100-soluble Aβ levels in hippocampal homogenates.

In 16-month-old 3xTg mice, numerous MXO4^+^ plaques were found in the *subiculum* and CA1 region of the hippocampus (**Fig. 3a**) and fewer Aβ42^+^ dense core plaques (not shown). Modulation of reactive astrocytes by SOCS3 or JAK2ca had no detectable impact on hippocampal amyloid plaques stained by MXO4 (**Fig. 3a**) or Aβ42 antibody (**Supplemental Fig. 3**). The number of plaques and their average individual size were identical between groups (**Fig. 3b, c and Supplemental Fig. 3**). Similarly, immunoreactivity in the *subiculum* and pyramidal layer of the CA1 region for intracellular Aβ labelled by the 4G8 antibody was identical between groups (**Supplemental Fig. 3**).

**Figure 3.**
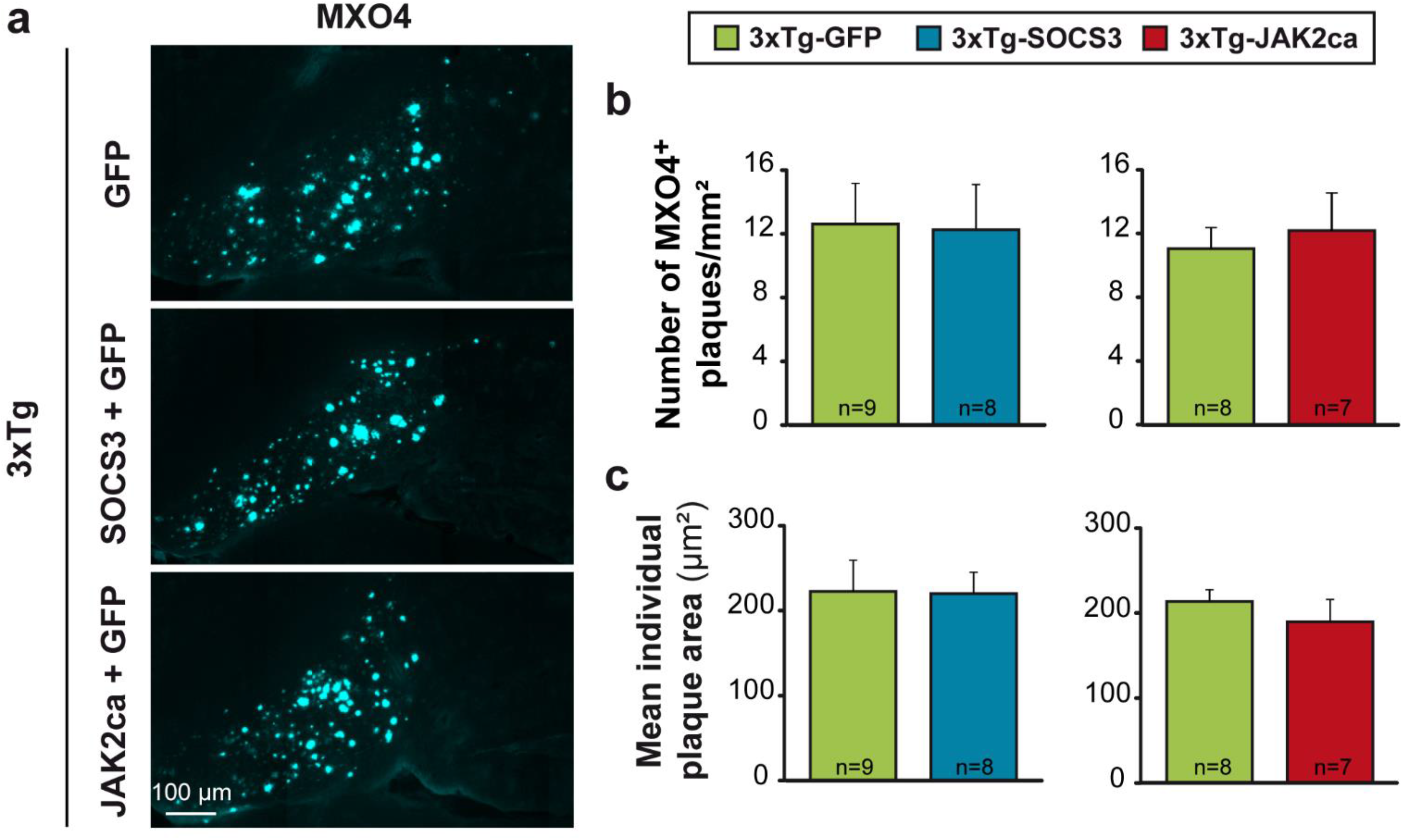
Modulation of reactive astrocytes does not impact amyloid load in 16-month-old 3xTg mice. **a**, Representative images of amyloid plaques labelled with MXO4 (cyan) in 16-month-old 3xTg-GFP, 3xTg-SOCS3 and 3xTg-JAK2ca mice. **b**, Quantification of the total number of MXO4^+^ plaques in the whole hippocampus shows no difference between groups. **c**, The mean size of individual MXO4^+^ amyloid plaques is also similar between groups. Mann-Whitney test.

We next measured Triton X-100 soluble Aβ levels by MSD® assays on protein homogenates prepared from the hippocampus of 3xTg-GFP, 3xTg-SOCS3 or 3xTg-JAK2ca mice. At the age of 16 months, only Aβ42 was reliably detected and quantified (Aβ40 was undetectable in 2 to 4 samples/group). Hippocampal Aβ42 concentrations were not significantly modified by SOCS3 or JAK2ca (**Fig. 4a**). Accordingly, expression of several proteins involved in APP metabolism were expressed at similar levels in 16-month-old in 3xTg-GFP, 3xTg-SOCS3 and 3xTg-JAK2ca mice [APP; BACE1, the β-secretase that cleaves APP through the amyloïdogenic pathway; insulin degrading enzyme (IDE) that degrades Aβ; and apolipoprotein E (ApoE) a protein secreted by astrocytes, which binds and clears cholesterol and Aβ (**Supplemental Fig. 4**)].

**Figure 4.**
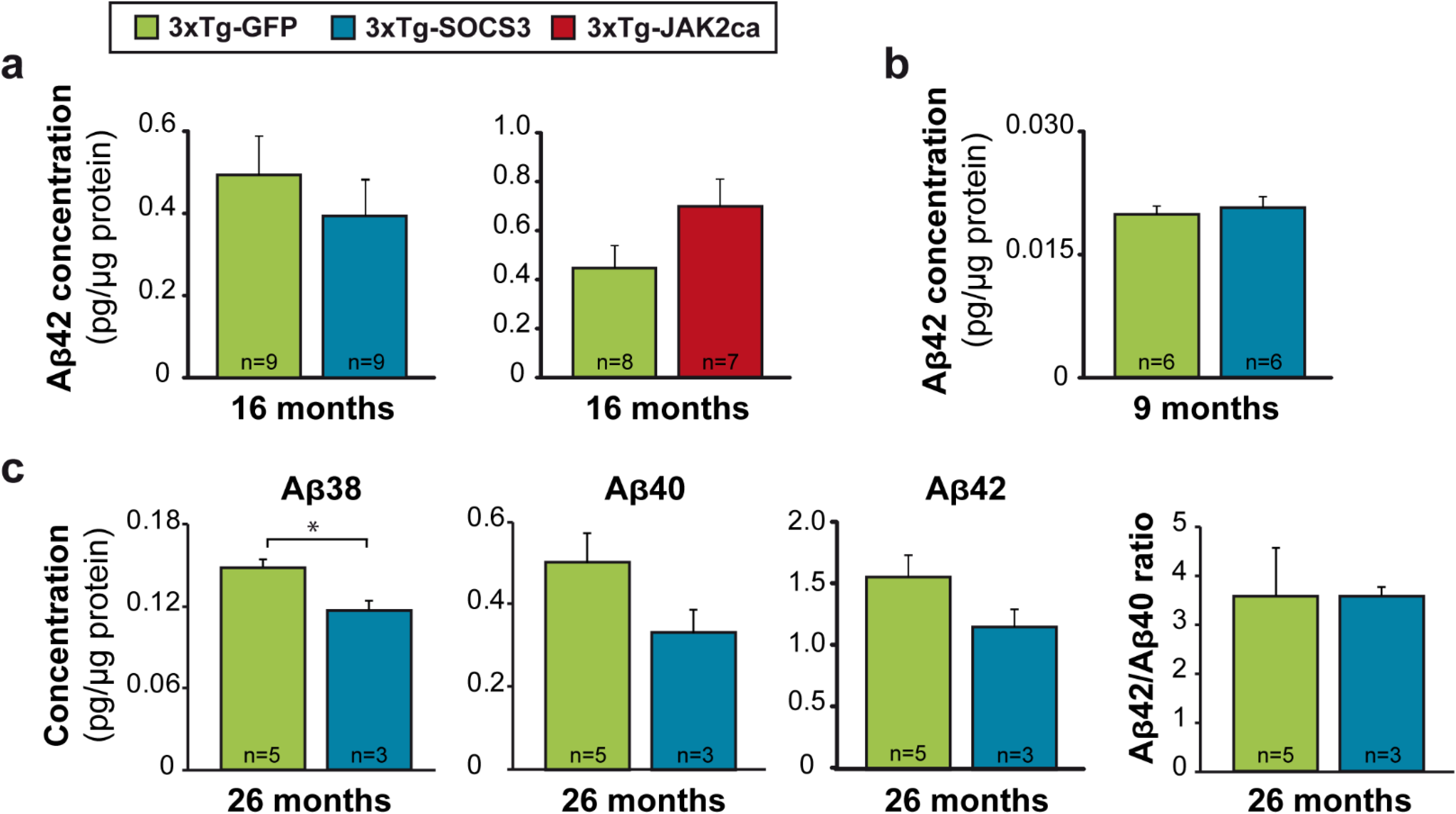
Modulation of reactive astrocytes does not impact soluble Aβ levels in 3xTg mice. MSD® measurement of Aβ42 concentration in Triton-X100 soluble protein homogenates prepared from the hippocampus of 16-month-old 3xTg-GFP, 3xTg-SOCS3 and 3xTg-JAK2ca mice (**a**), or in 3xTg-GFP and 3xTg-SOCS3 mice at the age of 9 (**b**) and 26 (**c**) months. Aβ42 levels increase with age, but are not changed by SOCS3 or JAK2ca in 3xTg mice. **c**, Aβ40 levels, Aβ42/40 ratio are similar in 3xTg-GFP and 3xTg-SOCS3 groups, while Aβ38 levels are significantly reduced by SOCS3 in end-stage 26-month-old 3xTg mice. * *p* < 0.05. Mann-Whitney test.

**Figure 5.**
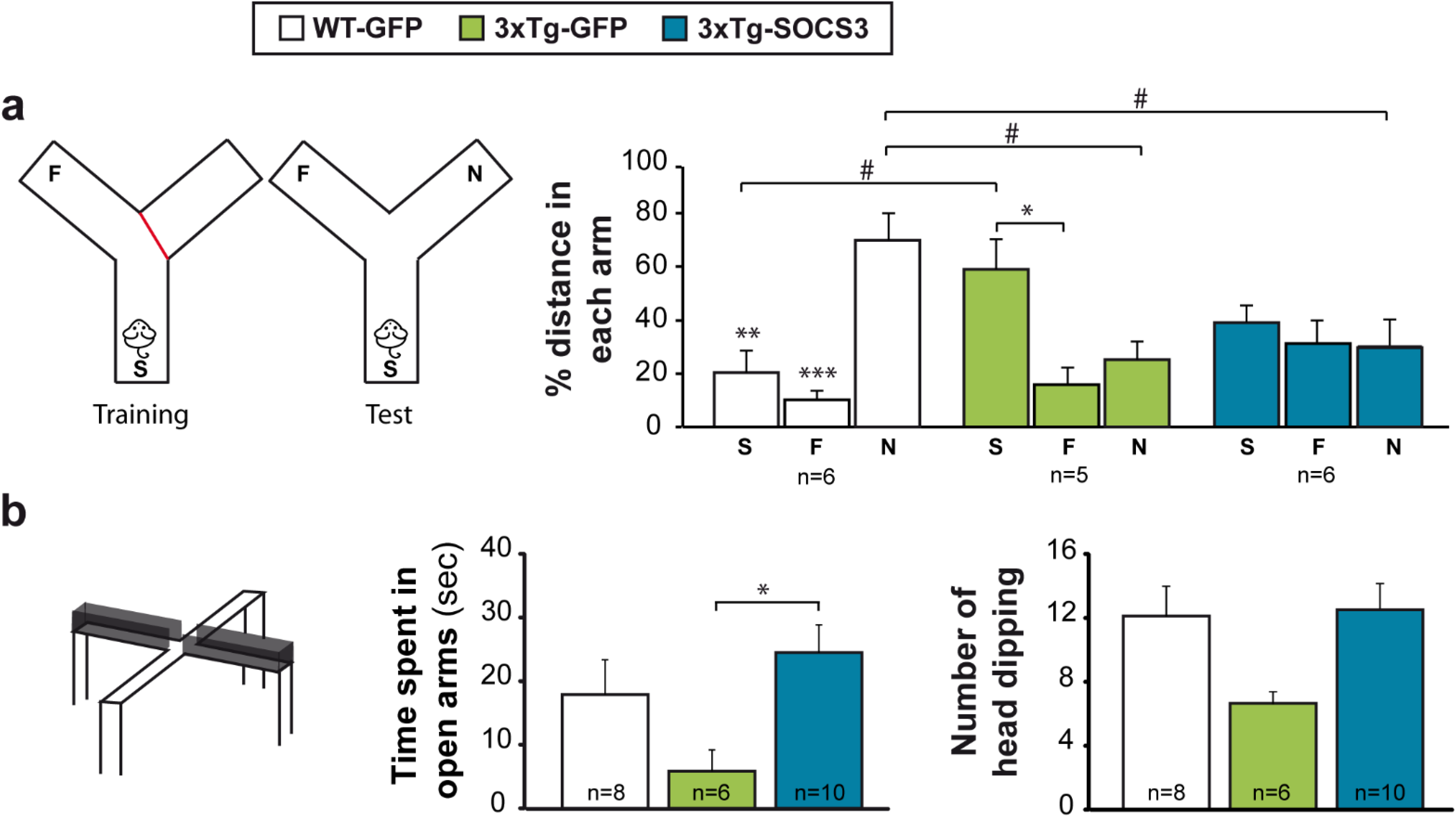
SOCS3-inbition of reactive astrocytes does not correct memory deficits but reduces anxiety in 3xTg mice. **a**, In the test phase of the Y maze, WT-GFP mice preferentially explore the novel arm (N) rather than the start (S) or the familiar arm (F). In contrast, 14-month-old 3xTg-GFP mice show no preference for the novel arm over the familiar arm, reflecting an altered working memory, and they significantly traveled more distance in the start arm than WT-GFP mice, suggesting increased anxiety. 3xTg-SOCS3 mice did not stay within the start arm and explored all arms similarly, suggesting that SOCS3 reduces mouse anxiety but does not correct short-term memory. * *p* < 0.05, ** *p* < 0.01, *** *p* < 0.001 (between arms within the same group), # *p* < 0.05 (between groups, for the same arm). Two-way (group, arm) ANOVA and Tuckey post-hoc test. **b**, In the elevated plus maze test, 3xTg-GFP mice tend to spend less time and perform less head dipping in open arms than WT-GFP mice. SOCS3 expression in hippocampal astrocytes significantly increases the time spent in open arms. * *p* < 0.05. ANOVA and Tuckey post-hoc test.

To study whether reactive astrocytes impact Aβ levels at an earlier or later stage of the disease, we performed the same measurement of Aβ levels in 9-month-old and 26-month-old female 3xTg-GFP and 3xTg-SOCS3 mice. Again, Aβ42 concentrations were identical between groups at both ages (**Fig. 4b, c**). Aβ38 and Aβ40 reached detectable levels in 26-month-old 3xTg mice. The Aβ42/Aβ40 ratio was not impacted by SOCS3 but concentrations of Aβ38, the least toxic form (Li et al., 2018), were significantly decreased by SOCS3 (**Fig. 4c**).

Overall, our analysis demonstrates that JAK2-STAT3-dependent reactive astrocytes do not overtly contribute to amyloid metabolism and deposition in 3xTg mice, at three different disease stages.

### 3.4. Inhibition of reactive astrocytes improves anxiety but not memory in 3xTg mice

We previously showed that hippocampal reactive astrocytes contribute to deficits in synaptic transmission and plasticity in 9-month-old 3xTg mice (Ceyzériat et al., 2018). Here, we analyzed whether synaptic restoration by SOCS3 translates into cognitive improvement in 3xTg mice.

We tested whether SOCS3-mediated inhibition of reactive astrocytes improved spatial memory deficits on the Y maze (Carroll et al., 2007) and anxiety on the elevated plus maze (Blanchard et al., 2010). WT-GFP, 3xTg-GFP mice and 3xTg-SOCS3 were first tested at 8 months. 3xTg mice only showed a tendency of altered behaviors at this early stage (data not shown), therefore mice were tested again at the age of 14 months.

Each mouse was placed on the Y maze to explore two arms (the start and familial arms, one being closed) for 15 min of training (**Fig. 5a**). After 5 min in their home cage, the mouse was placed again in the start arm of the maze with the three arms open. In average, WT-GFP, 3xTg-GFP and 3xTg-SOCS3 mice traveled the same distance during the test session, showing equivalent exploration activity (data not shown), but they explored the three arms differently (*p* < 0.0005 for group × arm effect, two-way ANOVA, **Fig. 5a**). WT-GFP mice displayed the expected preference for the novel arm during the test session. On the contrary, 3xTg-GFP mice displayed a significance preference for the start arm and explored similarly the novel and familiar arms. Such lack of interest for the novel arm reveals alteration in working memory, while the preference for the start arm suggests increased anxiety to explore more distant arms. SOCS3 did not improve working spatial memory in 3xTg mice, but it reduced their anxiety, as they did not remained in the start arm and explored the three arms similarly (**Fig. 5a**).

These mice were also tested on the elevated plus maze. The time spent in the open arms of the maze was used as a surrogate for the level of anxiety. 3xTg-GFP mice tended to spend less time and to perform less head dipping in open arms, an indication of higher anxiety. SOCS3 reduced anxiety in 3xTg mice, by significantly increasing the time spent in the stressful open arms (**Fig. 5b**).

Our data show that inhibition of reactive astrocytes by SOCS3 does not alleviate memory deficits in 3xTg mice but reduces anxiety levels.

## 4. DISCUSSION

Our multi-level preclinical analysis shows that modulation of hippocampal reactive astrocytes through the JAK2-STAT3 pathway does not significantly impact Tau or amyloid pathologies, at different stages of the disease in 3xTg-AD mice. However, SOCS3-mediated inhibition of reactive astrocytes reduced their anxiety on the elevated plus maze without restoring short-term spatial memory.

### 4.1. The JAK2-STAT3 pathway and reactive astrocytes

This study further demonstrates the central role played by the JAK2-STAT3 pathway in the control of reactive astrocytes, previously reported in other mouse models of neurodegenerative diseases (Ben Haim et al., 2015b; Ceyzériat et al., 2018; Reichenbach et al., 2019) and multiple diseases like ischemia, spinal cord injury or trauma (see, Ceyzériat et al., 2016 for review). Strikingly, we found that expression of SOCS3 or JAK2ca in astrocytes was able to respectively inhibit or exacerbate GFAP overexpression and morphological changes in hippocampal reactive astrocytes of 3xTg mice. These effects are extremely stable (observed after > 10 months post-injection) and allow analysis of reactive astrocytes at different disease stages. Previously, we showed that besides these two hallmarks, SOCS3 was also able to normalize the transcriptome of reactive astrocytes in neurodegenerative disease models (Ben Haim et al., 2015b; Ceyzériat et al., 2018). Similar reduction of reactive gene expression was found with the conditional knock-out of STAT3 in astrocytes in APP/PS1dE9 mice (Reichenbach et al., 2019) and mice with spinal cord injury (Anderson et al., 2016). Thus, activation of the JAK2-STAT3 pathway is necessary for both the induction and persistence of several molecular and morphological features of reactive astrocytes, identifying this cascade as a master regulator of astrocyte reaction in disease.

### 4.2. Reactive astrocytes do not influence molecular hallmarks of AD in 3xTg mice

We show that JAK2-STAT3-dependent reactive astrocytes do not significantly impact Tau and amyloid pathologies. This was found at three different stages of the disease (9, 16 and 24 months), ruling out stage-dependent effects. We only found a significant reduction in Aβ38 levels in the hippocampus of end-stage 3xTg-SOCS3 mice. Aβ40 and Aβ42 tended to be reduced in this group as well, but it did not reach significance. Aβ38 was shown to be less synaptotoxic than Aβ40 or Aβ42 (Li et al., 2018), suggesting that such an end-stage and marginal reduction in Aβ38 levels may not significantly alleviate neuronal dysfunctions. Therefore, JAK2-STAT3-dependent reactive astrocytes have only a minor role in amyloid processing in this model.

The lack of SOCS3 effects on amyloid deposition is in contrast with our previous study reporting significantly reduced number of MXO4^+^ amyloid plaques in APP/PS1dE9 mice with SOCS3 (Ceyzériat et al., 2018). This result is also in opposition with two studies reporting lower amyloid deposition after inhibition of reactive astrocytes through the calcineurin-NFAT pathway (Furman et al., 2012; Sompol et al., 2017) or after STAT3 conditional knockout in astrocytes of APP/PS1dE9 mice (Reichenbach et al., 2019). The fact that the same experimental manipulation of reactive astrocytes (through the JAK2-STAT3 pathway), in two AD mouse models, have different effects on the same disease outcome (amyloid plaques), assessed with the same method (MXO4) underlines that astrocyte reaction is more complex than expected. How can the discrepancies between these two AD models be explained? In 16-month-old 3xTg mice, both amyloid and Tau pathologies are present whereas only amyloid pathology occurs in APP/PS1dE9 mice. This important difference (in addition to other differences such as sex, genetic background, and genetic constructs) could change mechanisms of amyloid production, accumulation, aggregation and clearance. In addition, mutant Tau itself impacts astrocyte physiology (Dabir et al., 2006; Leyns and Holtzman, 2017) and could directly change their ability to clear amyloid.

But most importantly, accumulating evidence shows that astrocytes and their reactive counterparts are heterogeneous. Reactive astrocytes may adopt different molecular and functional states depending on the pathological context, and they could subsequently influence disease processes differentially (Anderson et al., 2014; Escartin et al., 2019). Therefore, even if the cause(s) for SOCS3 differential impact on amyloid deposition in 3xTg and APP/PS1dE9 mice were not identified, our studies illustrate that reactive astrocytes have complex and subtle effects *in vivo*, and call for caution in generalizing their roles in disease.

### 4.3. Inhibition of reactive astrocytes does not restore spatial memory but improves anxiety in 3xTg mice

We previously reported that SOCS3 expression in astrocytes fully restored early synaptic and LTP alterations in 9-month-old 3xTg mice (Ceyzériat et al., 2018). Here, we explored whether this normalization of synaptic activity translated into improved short-term memory and anxiety in 3xTg mice. We found that 8-month-old 3xTg mice do not display significant alterations on the Y maze and the elevated plus maze (data not shown). However, at 14 months, 3xTg mice displayed altered spatial memory and increased anxiety. Unexpectedly, blocking reactive astrocytes with SOCS3 improved anxiety but not working memory.

The absence of SOCS3 effect on working memory could be explained by the fact this type of memory involves other brain structures besides the targeted hippocampus. For example, it is now admitted that the prefrontal cortex is involved in spatial information processing (Funahashi, 2017). Accordingly, we found that SOCS3 was able to improve spatial learning but not memory retrieval on the Morris water maze in APP/PS1dE9 mice (Ceyzériat et al., 2018), further stressing the role of complex brain circuits for memory tasks. A strategy targeting the JAK-STAT3 pathway in a larger brain region through multiple injections, *i.v.* delivery of blood brain barrier permeant vectors encoding SOCS3 or pharmacological approaches would help uncover the effects of reactive astrocytes on memory impairment in AD mice.

Conversely, reduced anxiety in 3xTg-SOCS3 mice may be considered surprising, as AAV-SOCS3 was injected in the dorsal hippocampus, while anxiety-related behaviors are controlled by ventral hippocampus and amygdala (Bannerman et al., 2004; Calhoon and Tye, 2015). However, AAV may diffuse towards ventral hippocampus and gap junction-based astrocyte networks could transduce SOCS3 effects at distance from injected astrocytes, and regulate other hippocampal circuits controlling anxiety. Anxiety-related behaviors were evaluated on the elevated plus maze, as mouse propensity to explore anxiogenic open arms. Additional tests like the open field would be useful to better delineate anti-anxiogenic effects of SOCS3 in aged 3xTg mice. Interestingly, we observed that 3xTg mice remained in the start arm of the Y maze, an indication of increased anxiety and this was corrected as well by SOCS3, further demonstrating its positive effect on anxiety.

When do reactive astrocytes start to play a role in AD pathogenesis? Reactive glial cells (astrocytes and microglia) appear in response to an abnormal environment [e.g. dysfunctional neurons, presence of toxic oligomerized proteins (see, Ben Haim et al., 2015a, for review)]. But, positron emission tomography studies on prodromal AD patients showed that reactive glial cells may be present at early disease stages, before overt signs of amyloid deposition (Hamelin et al., 2016; Heneka et al., 2005). In accordance, our results indicate that reactive astrocytes contribute to early synaptic deficits, before amyloid deposition (Ceyzériat et al., 2018) and their inhibition may improve some behavioral outcomes without reducing Aβ and Tau pathologies.

## 5. CONCLUSIONS

Overall, the JAK2-STAT3 pathway appears as a core signaling cascade for reactive astrocytes. By modulating this pathway in different models of neurodegenerative diseases, we show that reactive astrocytes are multifaceted cells, with subtle functional effects, depending on the specific disease context. Our vector-based strategy is versatile and efficient to dissect out the roles of reactive astrocytes in different pathological conditions. A better understanding of the molecular changes occurring in reactive astrocytes and their impact on neurons is required to devise efficient therapeutic strategies targeting astrocytes.

## 6. FIGURE LEGENDS

**Supplemental figure 1.**
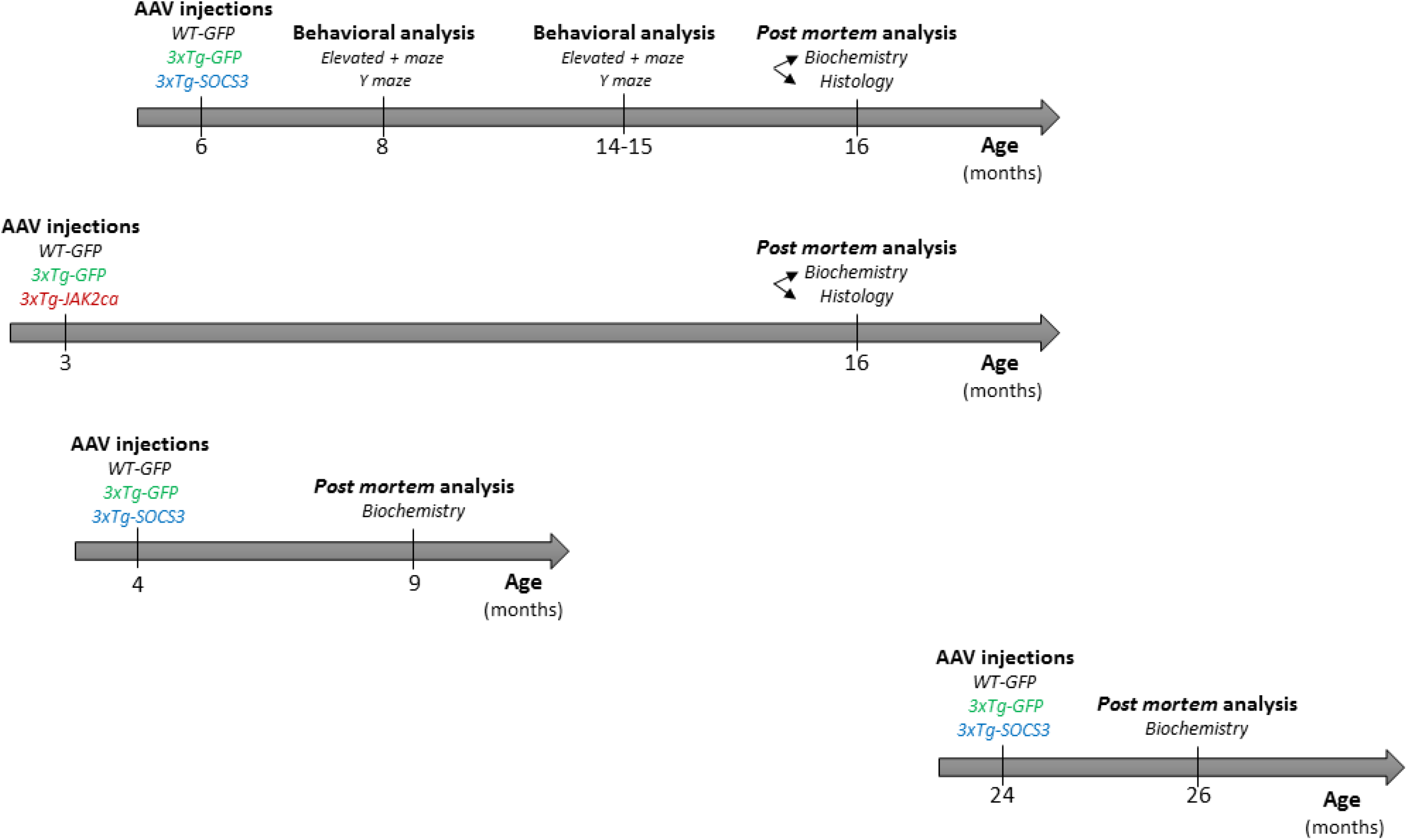
Experimental design. Four independent cohorts of female 3xTg mice and age- and gender-matched WT controls were injected and analyzed at the indicated ages.

**Supplemental figure 2.**
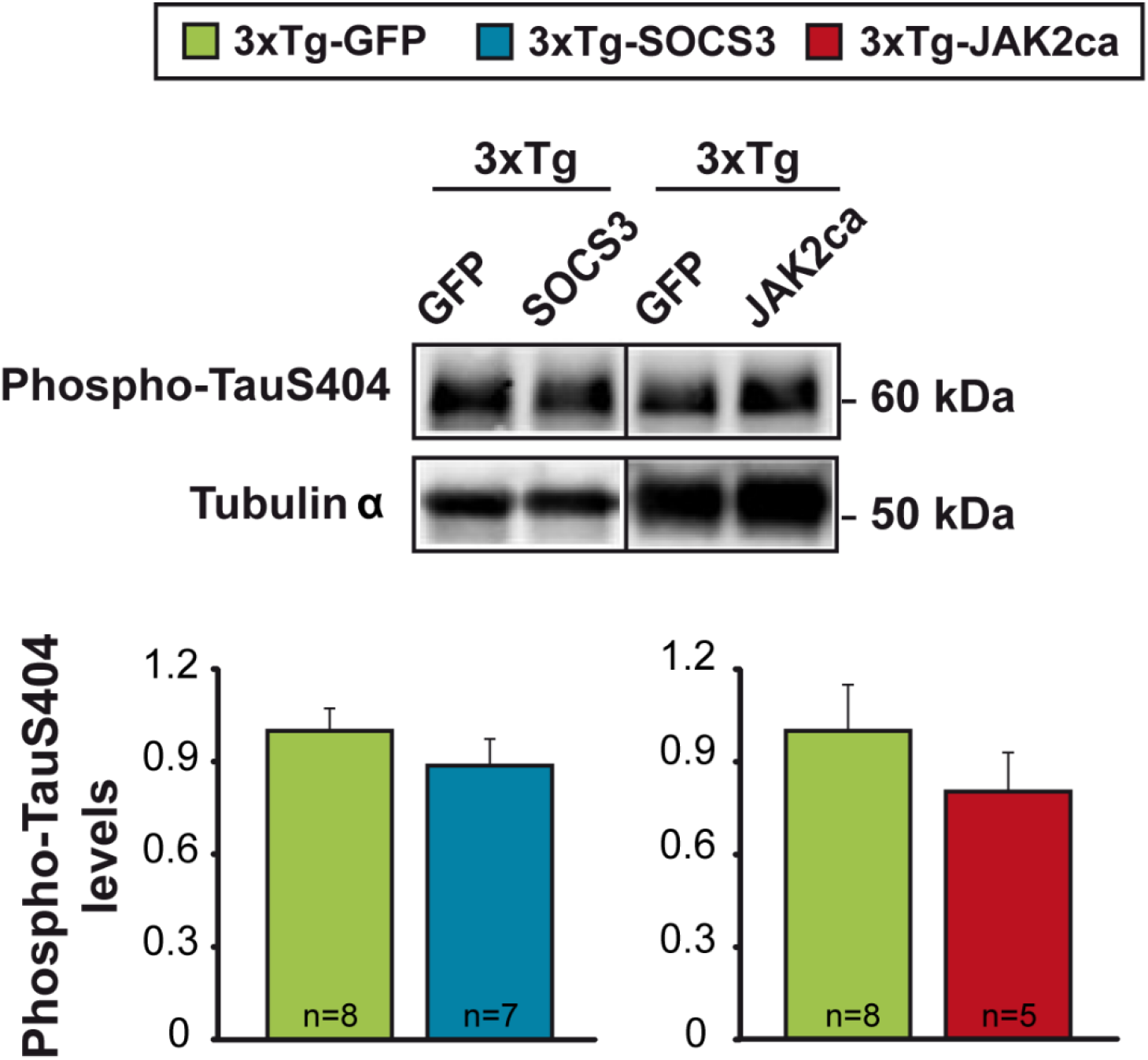
Modulation of reactive astrocytes does not impact phospho-Ser404-Tau levels in 3xTg mice. Representative western blotting and quantification of phospho-Ser404-Tau in 3xTg-GFP, 3xTg-SOCS3 and 3xTg-JAK2ca mice. Phospho-Ser404-Tau levels (normalized by tubulin α) are similar between groups. Mann-Whitney test. Quantifications are expressed relatively to the 3xTg-GFP group in each cohort, whose value was set to 1.

**Supplemental figure 3.**
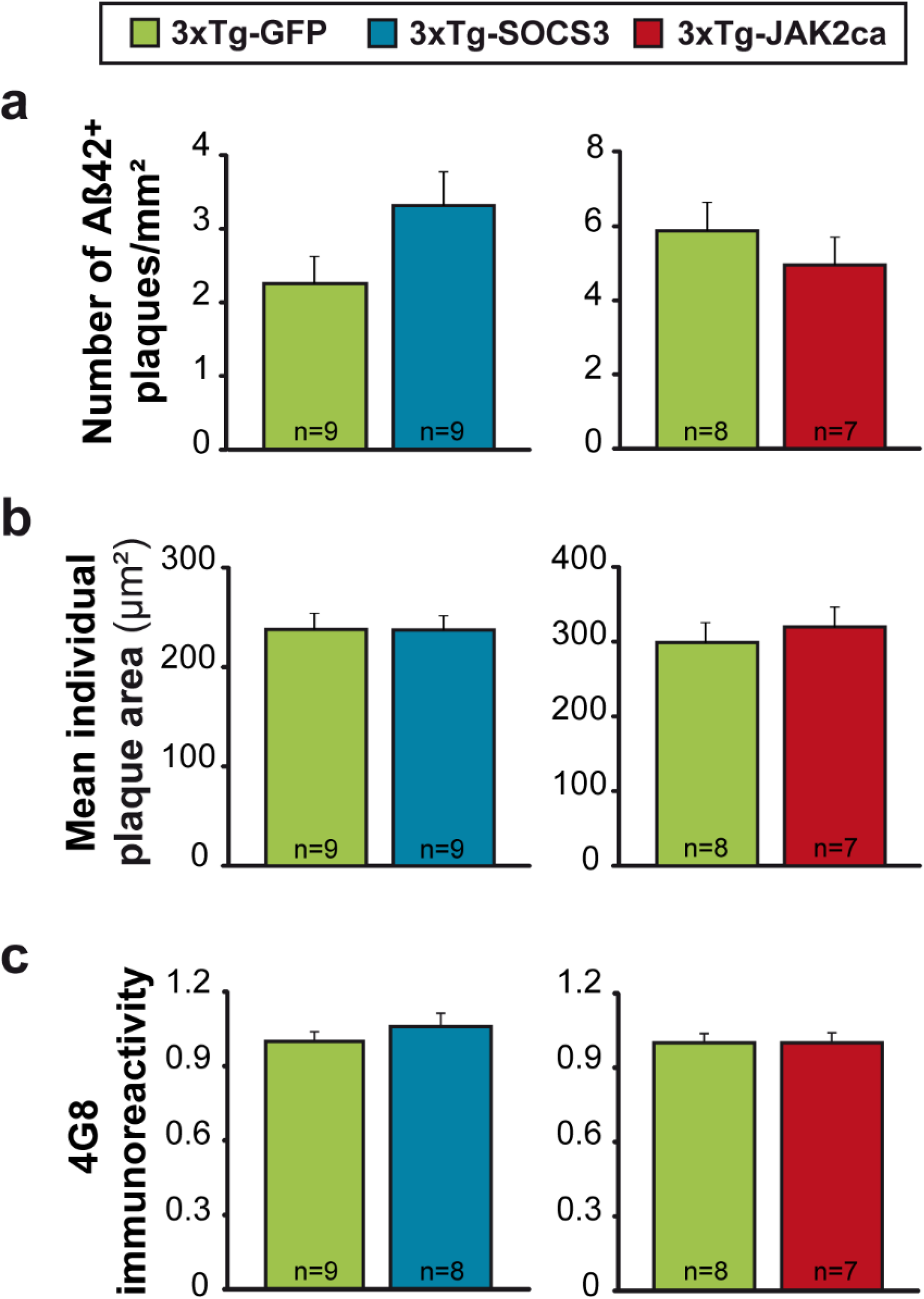
Modulation of reactive astrocytes does not impact amyloid pathology in 3xTg mice. **a, b**, Quantification of plaques labelled with an Aβ42 antibody in the whole hippocampus of 16-month-old 3xTg-GFP, 3xTg-SOCS3 and 3xTg-JAK2ca mice. The number (**a**) and average individual size (**b**) of Aβ42^+^ plaques are not impacted by SOCS3 or JAK2ca. N = 9/group. Mann-Whitney test. **c**, Quantification of 4G8 immunoreactivity in the *subiculum* and *stratum pyramidale* of 16-month-old 3xTg-GFP, 3xTg-SOCS3 and 3xTg-JAK2ca mice. There is no difference in 4G8 signal between groups. Mann-Whitney test. Quantifications in c are expressed relatively to the 3xTg-GFP group in each cohort, whose value was set to 1.

**Supplemental figure 4.**
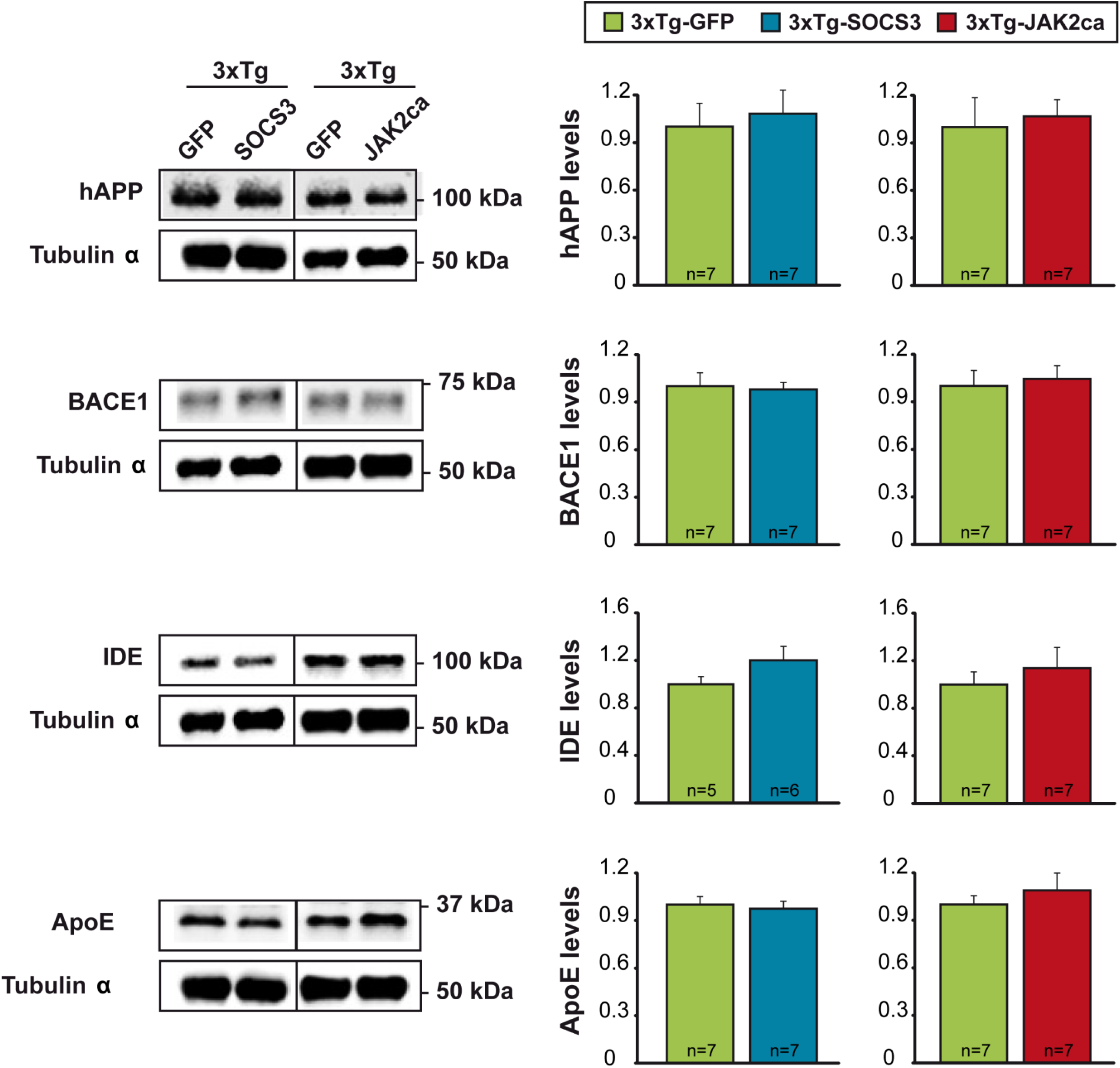
Modulation of reactive astrocytes does not impact expression of proteins involved in Aβ metabolism or clearance in 3xTg mice. **a**, Representative western blotting and quantification of human APP (hAPP) and key enzymes (BACE1, IDE) or transporter (ApoE) involved in Aβ production or clearance in 16-month-old 3xTg-GFP, 3xTg-SOCS3 and 3xTg-JAK2ca. Protein expression is similar between groups. Mann-Whitney test. Quantifications are expressed relatively to the 3xTg-GFP group in each cohort, whose value was set to 1.

## 7. Acknowledgements

We are grateful to D. Cheramy and Dr. E. Diguet (Servier) for sharing their expertise and equipment for MSD® kits. We thank A. Panatier and S.H.R. Oliet (NeuroCentre Magendie, Bordeaux) for providing 4-month-old 3xTg mice of the 3^rd^ cohort, as well as D. Gonzales, S. Laumond, J. Tessaire and people at the animal facility of the NeuroCentre Magendie for mouse care and genotyping. We thank L. Vincent, K. Bastide, P. Woodling and J.M. Hélies for their help with mouse housing and transfer.

